# “Ghost” fragment ions in structure and site-specific glycoproteomics analysis

**DOI:** 10.1101/2023.05.17.541150

**Authors:** Diana Campos, Michael Girgis, Qiang Yang, Guanghui Zong, Radoslav Goldman, Lai-Xi Wang, Miloslav Sanda

## Abstract

Mass spectrometry (MS) can unlock crucial insights into the intricate world of glycosylation analysis. Despite its immense potential, the qualitative and quantitative analysis of isobaric glycopeptide structures remains one of the most daunting hurdles in the field of glycoproteomics. The ability to distinguish between these complex glycan structures poses a significant challenge, hindering our ability to accurately measure and understand the role of glycoproteins in biological systems. A few recent publications described the use of collision energy (CE) modulation to improve structural elucidation, especially for qualitative purposes. Different linkages of glycan units usually demonstrate different stabilities under CID/HCD fragmentation conditions. Fragmentation of the glycan moiety produces low molecular weight ions (oxonium ions) that can serve as a structure-specific signature for specific glycan moieties, however, specificity of these fragments has never been examined closely. Here, we investigated fragmentation specificity using synthetic stable isotope-labelled glycopeptide standards. These standards were isotopically labelled at the reducing terminal GlcNAc, which allowed us to resolve fragments produced by oligomannose core moiety and fragments generated from outer antennary structures. Our research identified the potential for false positive structure assignments due to the occurrence of “Ghost” fragments resulting from single glyco unit rearrangement or mannose core fragmentation within the collision cell. To mitigate this issue, we have established a minimal intensity threshold for these fragments to prevent the misidentification of structure-specific fragments in glycoproteomics analysis. Our findings provide a crucial step forward in the quest for more accurate and reliable glycoproteomics measurements.

**Graphical abstract:** 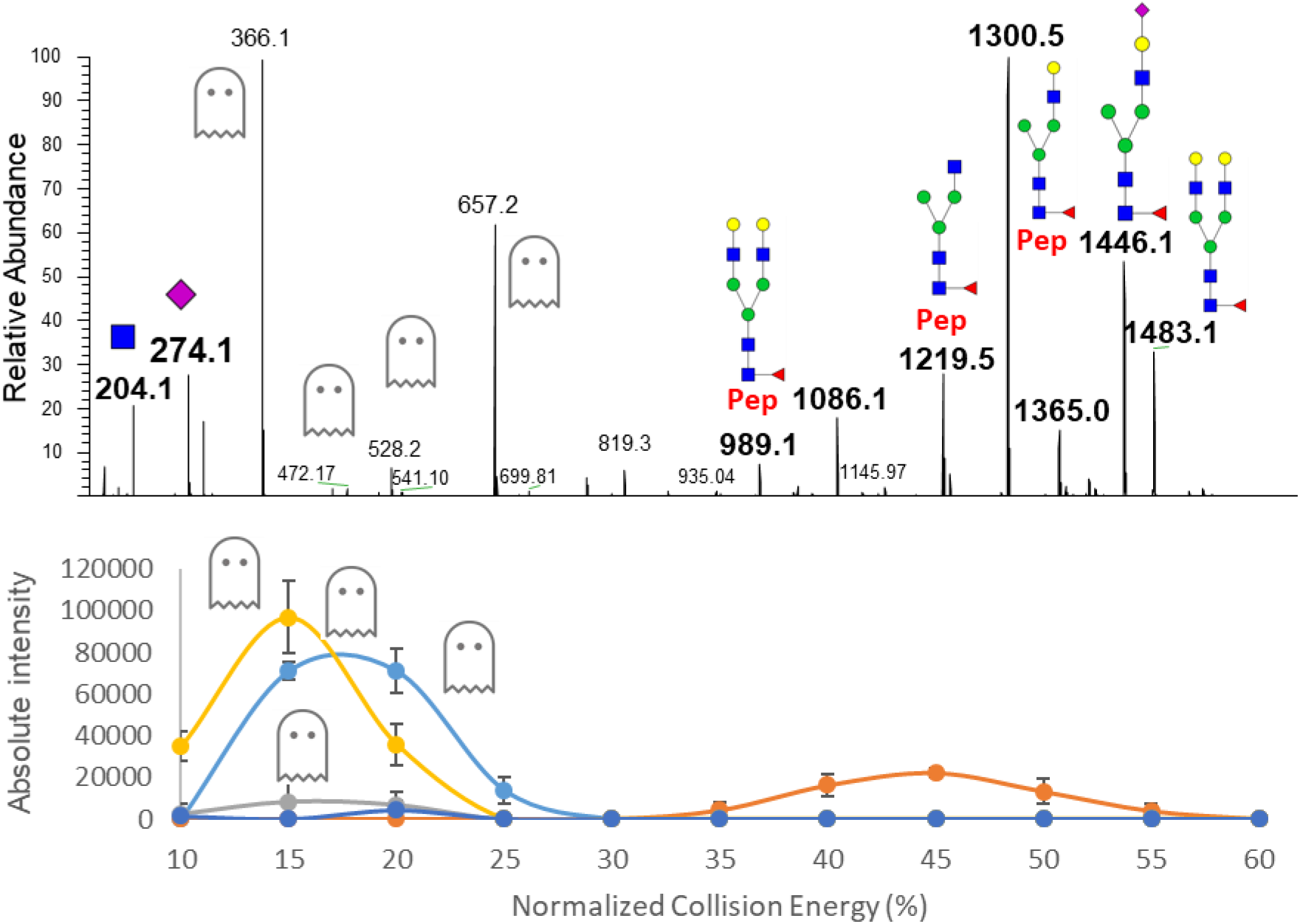

## LETTER

Precise characterization of protein glycosylation, in particular glycan structure analysis at a peptide specific site, has always been hampered by the microheterogeneity and nonlinear structural complexity of glycans ^1^. In an attempt to overcome this challenge, mass spectrometry (MS) workflows and glycoproteomics software packages have been heavily implemented ^2, 3^. Partial characterization of glycan composition, intact precursors or fragment ions, were analysed by a variety of software tools such as Byonic ^4^, pGlyco 2.0 ^5^, GPQuest ^6^, GPSeeker, MetaMorpheus ^7^, MSFragger-Glyco ^8^, and StrucGP ^9^. Although these types of software can serve as valuable tools for structural and site-specific glycan interpretation, this analysis still requires a good degree of manual data examination/curation, as the software readouts may lead to the assignment of false positive oxonium ion fragments.

Researchers have made noteworthy discoveries regarding glycan rearrangements during collision events at specific energies. These observations include rearrangements of hexose, as demonstrated by Wuhrer, Koeleman and colleagues in 2009 ^10^, and fucose, as shown in studies by Wuhrer, Koeleman et al. in 2006 ^11^ and Acs, Ozohanics et al. in 2018 ^12^. The results of these studies emphasized the critical importance of setting precise thresholds for particular fragments. This practice is crucial to prevent the erroneous identification of motifs that are unique to a structure and could potentially cause significant problems. Our team, along with other researchers, have developed a technique to determine the structural information from the tandem mass spectra of glycopeptides ^13, 14^. We used low collision energy beam type fragmentation to precisely assign outer antennary structural motifs and, in addition, we used structure specific fragments for their quantitation ^15, 16^. By modulating the collision energy, we are able to identify specific glycopeptide motifs with unparalleled accuracy (Figure 1). However, our work with synthetic standards has revealed a discrepancy between the structure-specific ions and their synthetic counterparts.

**Figure 1.**
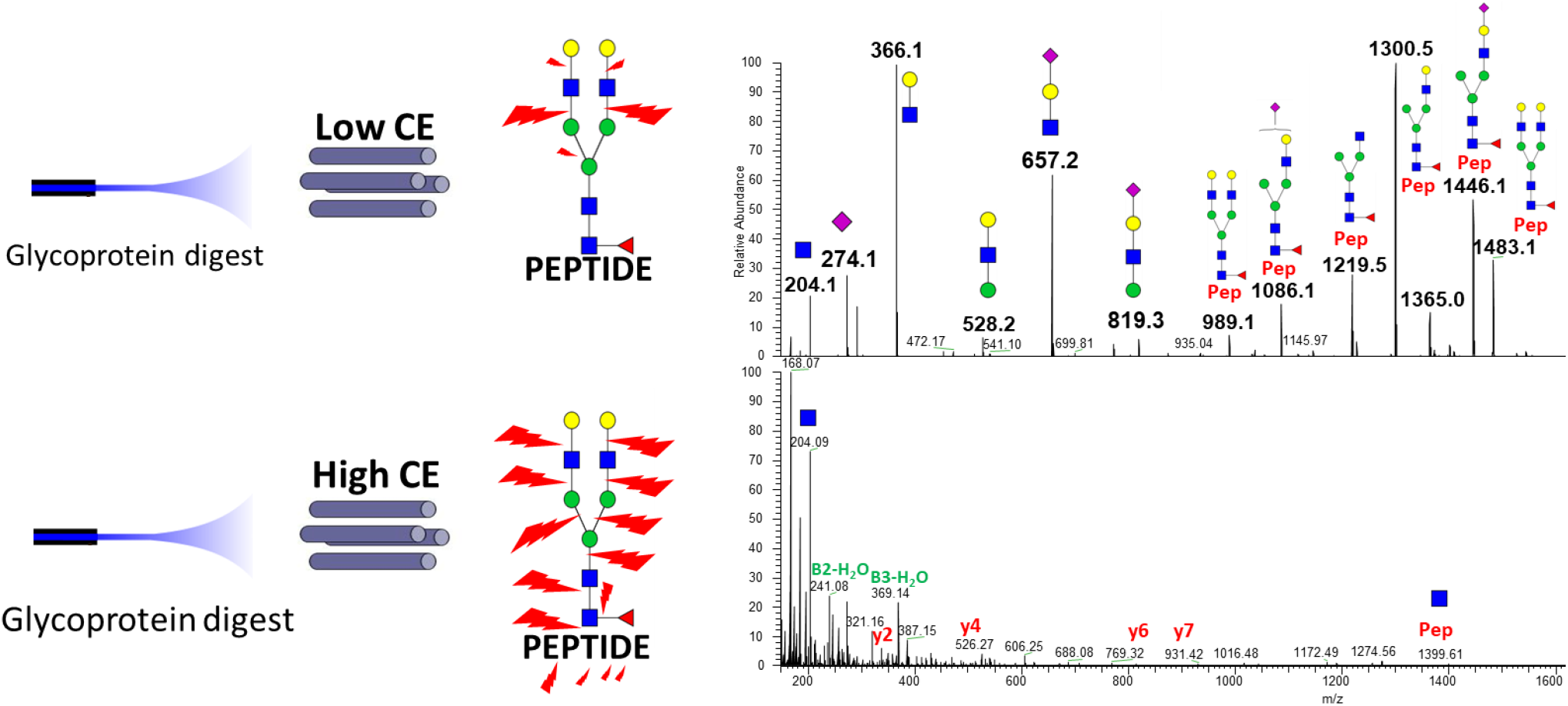
MS/MS spectra of the following structural motifs of G2FS IgG glycopeptide recorded under low collision energy NCE 10 (upper panel) and high collision energy NCE 45 (lower panel).

In this study, we utilized synthetic isotope-labelled standards (SIS) of glycopeptides and CE modulation to explore the existence of false positive signatures of specific outer antennary structures such as LacdiNAc, outer arm fucosylation, or outer arm GlcNAc sialylation and fucosylation, known as “Ghost “ fragment oxonium ions. To ensure the accuracy of our findings, we used IgG glycopeptide standards with ^13^C stable isotope incorporated into the reducing terminal GlcNAc, which creates a 6 Dalton difference compared to the natural IgG1 glycopeptides ^17^. This step was crucial in our research to avoid the contamination by natural isomers s that could compromise the reliability of our results.

Our LC-MS/MS-PRM workflow (detailed in the Supplementary information/methods) allowed us to record High Collision Dissociation (HCD) spectra of the tryptic digest of IgG1 glycopeptides and investigated fragmentation of five IgG glycopeptide standards (G0F, G1F, G2F, S1G2F, and S2G2F structures). We created separate PRM methods with transition lists with Normalized Collision Energy (NCE) range 10-60 using step 5 for each glycopeptide standard, and recorded spectra for charge state 3+ and 2+. This allowed us to estimate the relative abundance of “Ghost” oxonium ions to prevent false positive glyco-structure assignments.

For instance, fragment m/z 407.2 is commonly described and used as the signature for LacdiNAc (GalNAc-GlcNAc) outer antennae specific structure ^9^. However, isobaric diHexNAc fragment can theoretically be produced by fragmentation of the core of chitobiose (GlcNAc-GlcNA) (Figure 2). It is practically impossible to separate these isobaric ions in complex biological matrices without structure-specific separation of oxonium ions utilizing ion mobility mass spectrometry ^18^. Incorporating stable isotope ^13^C into GlcNAc produces a 6 Dalton difference between the “heavy” and “light” isotopic glycopeptides resulting in fragment ion m/z 413.2 in the diHexNAc fragment ion created by the chitobiose core fragmentation (Figure 2). Thus, with CE modulation, we were able to calculate a threshold for the false-positive LacdiNAc structure assignment (Figure 3 and Table 1). The area of the integrated peak was used for further data processing. We determined that the intensity of false positive LacdiNAc ion could be up to 3.58% of relative intensity at collision energy NCE 45. Fucose rearrangement was described previously by us and others ^11, 12, 14^. We evaluated the production of false-positive LacNAc-fucose ion (m/z 512.2), SialoLacNAc-fucose ion (m/z 803.3) and GlcNAc-fucose ion (m/z 350.1) and determined that the false outer arm LacNAc-fucose fragment could be up to 2.89%, the SialoLacNAc-fucose fragment up to 3.19%, and GlcNAc-fucose up to 1,81%, all at collision energy NCE 15. In case of the extended structures such as sialylated glycans the false production is whole fucosylated antenna (m/z 803.2) followed by fucosylated antenna fragments (m/z 512.2) and fucosylated GlcNAc (m/z 350.1) with the optimum slightly shifted to higher collision energies. Fragment m/z 350 is in this case specific for outer antennary structure because terminal GlcNAc is stable isotope labelled and fragment m/z 356 was not identified in any spectra. Therefore, false positive determination of outer antennary fucosylation is high under low collision energy commonly used for the glycan structure assignment.

**Table 1.** Threshold percentages of MS/MS relative abundance of “Ghost” oxonium ions for false positive glyco-structure assignment.

| *IgG santard* | *Fragment ion m/z* |  |  |  |  |
| --- | --- | --- | --- | --- | --- |
|  | **413.2 (407.2) (%)** | **350,1 (%)** | **512.2 (%)** | **803.3 (%)** | **495.2 (%)** |
| **G0F** | 0,7 (NCE 45) | 1,43 (NCE 15) | 0,48 (NCE 15) |  |  |
| **G1F** | 0,48 (NCE 45) | 1,81 (NCE 15) | 2,18 (NCE 15) | - | - |
| **G2F** | 0,41 (NCE 45) | 0,29 (NCE 15) | 2,89 (NCE 15) | - | - |
| **S1G2F** | 3,27 (NCE 45) | 0,2 (NCE 20) | 2,47 (NCE 15) | 1,08 (NCE 15) | 0,2 (NCE 15) |
| **S2G2F** | 3,58 (NCE 45) | 0,24 (NCE 20) | 1,67 (NCE 20) | 3,19 (NCE 15) | 0,58 (NCE 15) |

**Figure 2.**
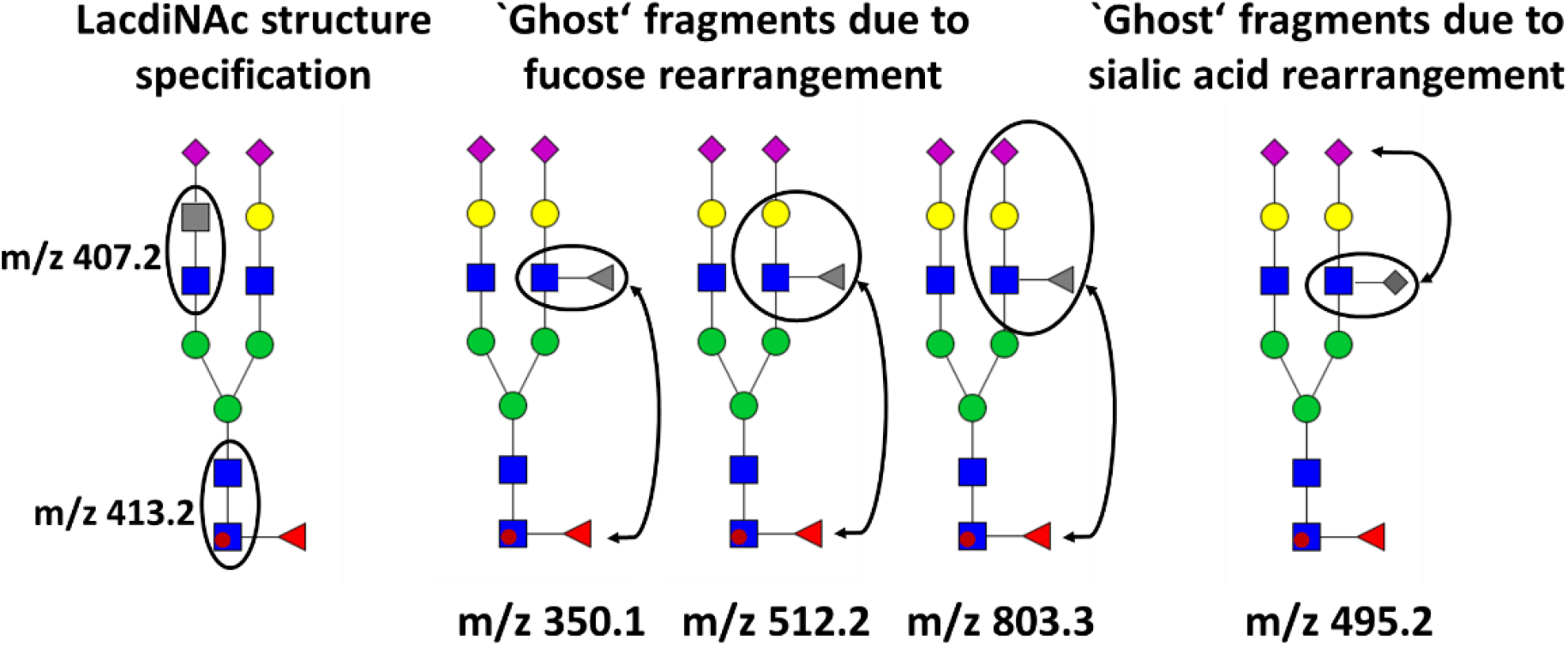
Schematic depiction of LacdiNAc specific ions and fucose and sialic acid rearrangements.

**Figure 3.**
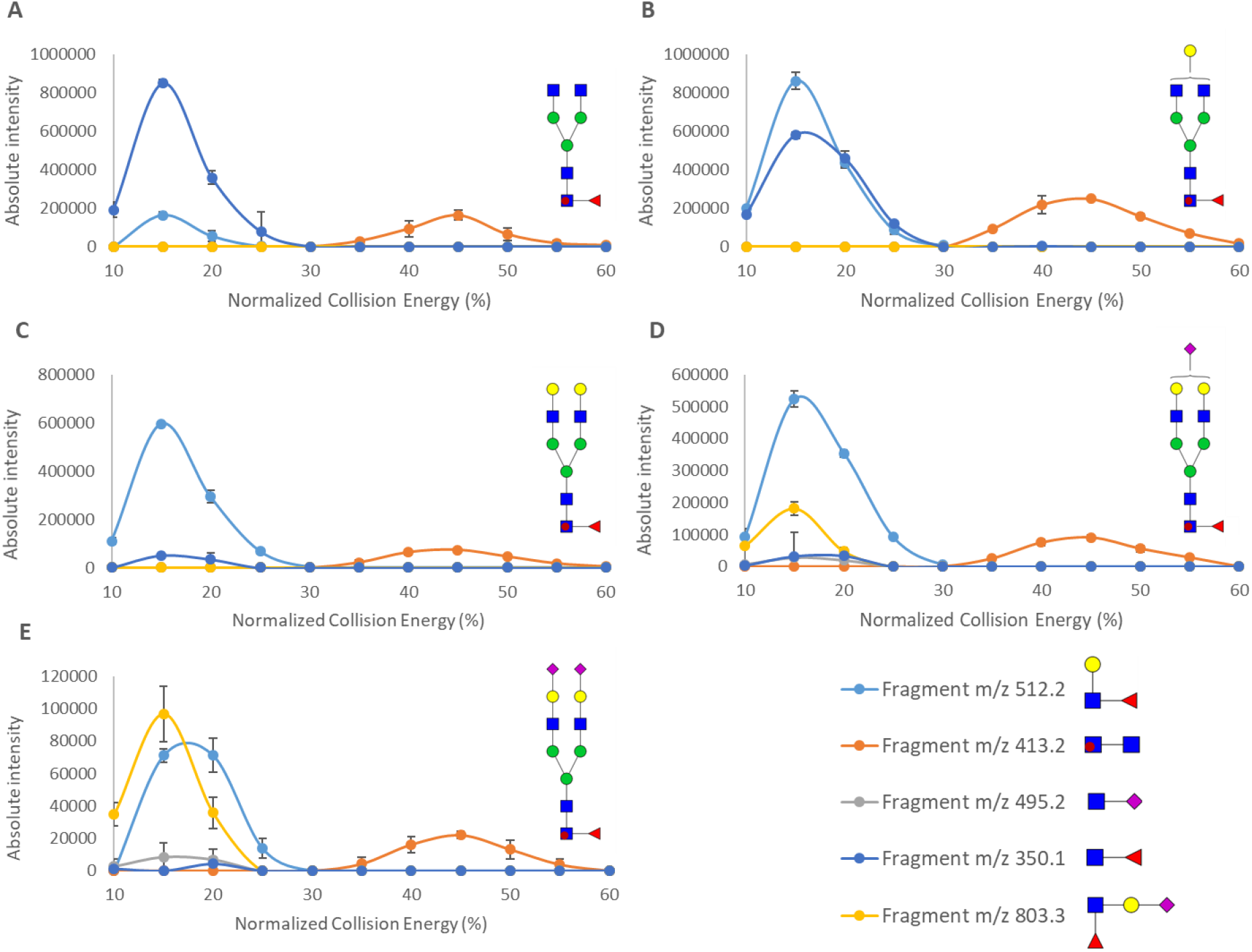
Collision energy settings for PRM of labelled IgG1 FC glycopeptide standards. Absolute intensity of fragment ions m/z 350; 413.2; 512.2; 495.2 and 803.3 as function of collision energy for structures G0F (A), G1F (B), G2F (C), SG2F (D) and S2G2F €.

The sialic acid attached to the GlcNAc is an atypical sialylated glycan that was described in human samples in some previous studies ^19, 20^. These glycoproteomics studies used ion (SA-HexNAc) m/z 495.2 as the signature for this unusual (non-human) structure. The m/z 495.2 ion is commonly produced by fragmentation of a sialo-LacdiNAc structure (Figure 2) or Sialo-T antigen^14^.To our knowledge, this is the first-time sialic acid rearrangement was described, similar to fucose rearrangement using HCD fragmentation. Production of m/z 495.2 ion showed very similar characteristics compared to fucose rearrangement and could be falsely used for the determination of SA-GlcNAc linkage. We determined a threshold percentage of 0.58% of 495.2 at 15% collision energy relative intensity as the highest false positive signal of SA-HexNAc outer arm structure. As expected, CE dependent fragmentation, profiles for fucose and sialic acid rearrangement demonstrated similar pattern (Figure 3). In other words, if the percentage of this m/z 495.2 ion exceeds 0.58% under the specified conditions, it is likely to be a false positive result for the SA-GlcNAc linkage. Additionally, the fragmentation profiles for fucose and sialic acid rearrangement showed a similar pattern. This means that the sialic acid rearrangement can be distinguished from fucose rearrangement based on the different patterns observed in the fragmentation profiles. Overall, the discovery of sialic acid rearrangement is an important contribution to the field of glycobiology and has implications for the accurate identification of specific glycan structures in human samples.

Despite recent advances in glycopeptide analysis, there is still a lack of optimal fragmentation conditions to distinguish between false-positive fragments resulting from rearrangement and those produced by fragmentation of the mannose core. As a result, it is essential to establish threshold settings and conduct further investigations to minimize false assignments of glycopeptide structures. By doing so, we can improve the accuracy and reliability of glycopeptide analysis, ultimately leading to a better understanding of the roles of glycosylation in various biological processes.

## ACKNOWLEDGMENTS

We thank Ms. He Chen and Mr. Zhihao Zheng for technical assistance. This work was supported in part by the National Institutes of Health NIH grants U01CA230692 to MS

## Notes

### Competing Interest Statement

The authors have declared no competing interest.

