## Supplemental material for "“Ghost” fragment ions in structure and site-specific glycoproteomics analysis"

^a^Max-Planck-Institut fuer Herz- und Lungenforschung, Ludwigstrasse 43, Bad Nauheim, 61231, Germany; ^b^Department of Bioengineering, College of Engineering and Computing, George Mason University, Fairfax, VA, USA; ^c^GlycoT Therapeutics, College Park, MD, USA, ^d^Department of Chemistry and Biochemistry, University of Maryland, College Park, MD 20742; ^e^Department of Oncology, Lombardi Comprehensive Cancer Center, Georgetown University, Washington, D.C. ^f^Clinical and Translational Glycoscience Research Center, Georgetown University, Washington, D.C., 20057

**Contents:**

1. Materials and Methods
2. References

**MATERIALS AND METHODS**

**Synthesis of 13C-Labeled IgG1-Fc Glycopeptides**

IgG1-Fc glycopeptides were synthesized and purified as previously published by us ^1, 2^. After lyophilization, the synthesized glycopeptides were weighed on an accurate balance and further quantitated by analytic HPLC. In the following procedures, we used the IgG1 glycopeptide labelled standars with G1F, G2F, SG2F and S2G2F structures with extended sequence AKTKPREEQYNSTYRVVS.

**Glycopeptide Analysis by a Nano-LC-MS/MS-PRM workflow**

2 µg of each IgG1 glycopeptide stable Isotope labeled standards was dissolved in 20µl of 50mM AmBic buffer. The glycopeptide mixture was reduced by 5mM DTT at 56°C and alkylated by 15mM IAA for 30min. Peptide mixture was digested by trypsin in a ratio 1:50 at 37°C overnight. Glycopeptides were separated by capillary reverse phase (C18) nano-chromatography on in-house packed silica emitter tip (15cm x 75 µm) using Easy Nano chromatographic system (Thermo). Glycopeptides were separated at 0.3 μL/min as follows: starting conditions 2% ACN, 0.1% formic acid; 0−35 min, 2− 50% ACN, 0.1% formic acid; 35−42 min, 50−90% ACN, 0.1% formic acid; 42−47 min 90% ACN, 0.1% formic acid followed by equilibration to starting conditions for additional 3 min. Q-exactive HF mass spectrometer (Thermo) was used for MS and MS/MS analysis. Fragmentation spectra were recorded at 30 000 resolving power. 100 pg of tryptic glycopeptide mixture containing G1F, G2F, SG2F and S2G2F structures was injected on column. Separated Parallel

Reaction Monitoring (PRM) methods with transition lists with Normalized collision energy (NCE) range 10-60 using 5stpes were created for each glycopeptide standard. Two charge states (+2 and +3) were subjected to fragmentation analysis. Qual-browser software was used for qualitative and quantitative data processing. 3 replicates of each standard were measured.

**Glycopeptide Analysis by a Nano-LC-MS/MS-PRM workflow**

Xcalibur (Thermo) software was used for quantitative data processing. Processing methods were created for ion extraction from each PRM transition. Area of integrated peak was used for further data processing. Further data processing and graphing was carried out in Microsoft Excel.
